# Dynamic responses of human lung innate and adaptive immune cells highlight the roles of genes at asthma risk loci

**DOI:** 10.1101/2024.09.20.614132

**Authors:** Funing Tian, Donna C. Decker, Anne I Sperling, Nathan Schoettler

**Affiliations:** Department of Medicine, Section of Pulmonary and Critical Care, University of Chicago, Chicago, Illinois; Department of Medicine, Pulmonary and Critical Care Division, University of Virginia, Charlottesville, Virginia

**Keywords:** Lung resident immune cells, T cells, B cells, macrophages, asthma, non-classical HLA

## Abstract

**Rationale:** The lung is a unique immunological niche with diverse immune cell types. The effects of stimulation through innate and adaptive immune receptors on human lung immune cells has largely been extrapolated from studies of blood immune cells. While multiple immune cell types and many genes have been implicated as contributing to asthma, the dynamics of these in human lung immune cells following activation will yield insights into asthma pathogenesis and lung immunity more broadly.

**Methods and Measurements:** Human lung immune cells from 6 donors were isolated. Mixed leukocytes were treated separately with lipopolysaccharide (LPS), F(ab’)^2^-anti-human-IgM/IgG + IL4 and anti-CD3/CD28 for 4 and 18 hours and underwent single cell RNA sequencing (scRNAseq). Lung immune cell types were annotated, and gene expression compared across conditions. Genes at prior asthma-associated genetic loci were characterized across cell types, treatments and timepoints. Expression of non-classical class II genes associated with asthma, *HLA-DQA2* and *HLA-DQB2*, and their protein products was characterized with immunohistochemistry.

**Main Results:** We characterized gene expression in 116,697 lung immune cells. Cell-, treatment-, and timepoint-specific effects on gene expression were detected in all lung immune cell populations. Correlation of gene expression between lung and blood lymphocyte populations decreased following stimulation. Among the genes that were differentially expressed, 97 receptor:ligand pairs had changes with treatments. 96.0% of genes at asthma risk loci demonstrated differential expression in at least one cell type and at least one treatment. B cells were the cell type with the highest expression of *HLA-DQA2* and *HLA-DQB2* which increased with anti-IgM/IgG treatment and the HLA-DQβ2 protein was identified in lung B cells from a donor with asthma.

**Conclusions:** Human lung immune activation elicits a broad range of cellular responses that deviate from those of blood immune cells and are relevant to asthma. Lung B cells expressing *HLA-DQA2* and *HLA-DQB2* appear to be involved in a novel antigen presentation pathway that contributes to asthma risk.

## INTRODUCTION

The lung has constant exposures to air and inhaled materials and is a unique immunological niche and has frontline exposure to microbes. Lung immunity is mediated by a diverse cast of cells and includes both innate immune cell types such as macrophages and adaptive immune cells that include B cell and T cell populations.

Single cell RNA sequencing studies have provided detailed resolution of immune cells in tissues, including the lung(1–9), and have identified multiple subsets of each of these immune cell populations in human lungs. While some studies have evaluated lung immune populations following immune activation or viral infection, the majority capture a single snapshot of the immune landscape and cell states. More is known about blood immune cells following stimulation. For example, detailed responses of 17 different blood T cell subsets has been characterized following anti-CD3/CD28 treatment(10), and in another study the responses of 32 blood immune cell populations under resting and stimulated conditions has been evaluated(11). However, the blood does not have appreciable levels of immune cell types that are abundant in the human lung such as lung-resident alveolar macrophages or lung-resident T cells, and comprehensive assessment of innate and adaptive immune stimulation of human lung immune cells has not been assessed.

Asthma is a complex immune-mediated disease with genetic and environmental risk factors such as early life rhinovirus infection. Genetic studies have consistently associated asthma with genomic regions that harbor genes with prominent roles in immune responses(12–15). The function of some of these genes and their protein products have been extensively characterized and even become the target of therapeutics, such as IL-4. However, not all patients respond to these therapies, and the gene or genes at most asthma-risk loci that facilitate risk are not known. Moreover, which lung immune cells and under what cell states these genes are expressed remains a gap in the understanding of asthma pathogenesis.

Here, we sought to determine how immune cells in human lungs respond to stimulation through innate and adaptive immune receptors. We used scRNAseq to identify direct and indirect responses separately for lipopolysaccharide (LPS), anti-IgG/IgM and anti-CD3/CD28 treatments to define cellular responses at two timepoints. We establish that these responses differ from the responses blood immune cells. We reveal the dynamics of lung immune cell responses. We further demonstrate that genes nominated as asthma risk genes in genetic studies(14) have broad cell type- and treatment-specific responses and identify a role for asthma-associated non-classical human leukocyte antigen (HLA) class II molecules(16) in antigen presentation in human lung B cells.

## METHODS

### Human tissue procurement

Human lungs were obtained from organ donors whose lungs were not used for transplantation through the Gift of Hope Regional Organ Bank of Illinois. Donors with more than ten pack-years of tobacco use were excluded from this study. Because samples from this study were from deceased donors, this research was deemed to be non-human subjects research by the Institutional Review Board at the University of Chicago.

### Leukocyte isolation

Cells were processed as previously described(17, 18). Briefly, lung tissue was perfused with sterile fetal bovine serum and phosphate-buffered saline instilled in the pulmonary arteries. The right lower lobe was minced and digested into a single cell suspension. Mononuclear cells were isolated using Ficoll-Paque Premium (Sigma-Aldrich, Saint Louis, MO; Catalog Number GE-17-5442-02) and cryopreserved.

### Cell processing and treatments

Mononuclear cells were thawed and incubated with DNase for 10 minutes followed by processing using Ficoll-Paque Premium to remove dead cells and debris. Cells were resuspended in RPMI supplemented with 10% FBS and separately treated with lipopolysaccharide (LPS; 5μg/ml; eBiosciences, Catalog Number 00-4976) or F(ab’)^2^-anti-Human IgM/IgG (10μg/ml; Invitrogen, Catalog Number 16-5099086) + IL4 (20ng/ml; Cell Sciences, Catalog Number CRI104B) which we henceforth refer to as anti-IgM/IgG. Cells were resuspended in CTS OpTmizer Cell Media (ThermoFisher, Catalog Number A1048501) for treatment with anti-CD3/CD28 Dynabeads (ThermoFisher, Catalog Number 11161D). One media control experiment was performed using CTS OpTmizer Cell Media for comparison with RPMI media control. Cells were rested for two hours at 37°C prior to treatments. Anti-CD3/CD28 Dynabeads were removed from the culture media at the end of the experimental timepoint. Samples underwent cell hashtagging with cell multiplexing oloigo labeling using a 3’CellPlex Kit (10XGenomics, Catalog Number PN-1000261). To ensure highly viable cells needed for scRNAseq, samples were stained with Apotracker (Biolegend, Catalog Number 427402). For the LPS treatments, gating on lymphocyte and macrophage cells was performed using forward- and side-scatter and Apotracker negative cells were sorted a Symphony S6 cell sorter (BD Biosciences). To enrich for lymphocytes and reduce the number of macrophages in the anti-CD3/CD28 and anti-IgM/IgG treatments, leukocytes were stained with anti-CD20 efluor 506 (Invitrogen,Catalog number 69-0209-42), anti-CD3-PE (BioLegend, Catalog number300441) and anti CD45-APC (BioLegend, Catalog number 304037) antibodies and sorted on Apotracker negative cells and CD45+CD3+CD20- for the anti-CD/CD28 experiments and CD45+CD20+CD3- for the anti-IgM/IgG experiments with an CD45+ also sorted at 10% of the sorted lymphocyte populations. were sorted and after sorting cells from the gate that included macrophages was added at one tenth the number of sorted lymphocytes. Hashtagged and sorted cells were combined at a 1:1 ratio and immediately processed for scRNAseq.

### Single cell gene expression

Cells were processed for scRNAseq on a Chromium X system with a Chromium GEM Chip (10XGenomics, Catalog Number 1000422). Gene expression and hashtag libraries were generated using a 3’ and hashtag library (10XGenomics, Catalog Number 1000686). Libraries were sequenced on NovaSeq-6000 and NovaSeq-X-Plus (Illumina) with 150bp paired-end reads. Samples were sequenced targeting a median read depth of 30,000 reads per cell.

### Quality control and integration of single-cell genomics data

scRNA-seq data were processed and aligned to GRCh38 human genome reference with CellRanger (10XGenomics, v6.1.2). The resulting gene expression profile calculated per cell barcode was analyzed using Seurat(19) (v5.1.0) in R. Low-quality cells were filtered using the parameters min.cells = 3, percent.mt <20, and min.features = 200. Cells were further excluded if they were detected as doublets by both “centroids” and “medoids” with DoubletDecon(20) (v1.1.6) in R. Prior to integration, gene expression of individual samples was normalized using the ‘NormalizeData’ function with default parameters.

Integration of samples was performed by selecting the top 1000 highly variable features for anchor identification using the “rpca” method in the ‘FindIntegrationAnchors’ function. This was followed by data scaling, principal component analysis (PCA) and cell clustering of the integrated dataset. Uniform manifold approximation and projection (UMAP) was applied for further dimensionality reduction and visualization. Pearson’s correlation coefficients of gene expression across cell types and conditions were calculated with the “stat_cor” function from the ggpubr (v0.6.0) in R. Annotation of cell clusters was performed using CellTypist(2) (v1.6.3) with the “Human_Lung_Atlas” model (v2) in python.

### Differential gene expression, gene ontology and cell-to-cell communication analysis

Differentially expressed genes (DEGs) were calculated by comparing 1) simulated with unstimulated conditions for each cell type, and 2) B cells expressing *HLA-DQA/DQB2* with those that do not express any of the two genes. These comparisons were performed with the “FindMarkers” function in Seurat using a MAST test. Genes were considered as significant differentially expressed if they were with log_2_ fold-change ≥ 0.25 and adjusted p-value (P_adj_) ≤0.05. The significant DEGs were used as input for Gene Ontology (GO) enrichment analysis by gprofiler2(4) (v0.2.3) in R. Only enriched GO terms with false-discover rate-(FDR-) adjusted p-values < 0.05 were plotted (Figure 3). Cell-cell interaction was inferred from the significant DEGs using CellChat (v.2.1.2, https://github.com/jinworks/CellChat) in R.

### Comparison of gene expression 1) in blood and lung immunes cells, and 2) for genes at asthma loci

Lung and blood immune cell gene expression at untreated and following immune stimulation were compared. Gene expression in blood immune cells was previously reported(11). Gene expression in this dataset was normalized with the function “normalize.quantiles” from preprocessCore (v1.66.0) in R. For lung immune cells, only the expression of significant DEGs under anti-IgM/IgG for B cells, anti-CD3/CD28 for CD4 T cells, CD8 T cells and NK/γδ cells and LPS for macrophages were considered for lung immune cells. Pearson’s correlation coefficients were calculated on the average gene expression of blood and lung immunes cells. Expression of asthmatic genes under treatments was compared with the unstimulated condition across cell types and was assessed for significant differences using Mann-Whitney U-tests in R.

### Immunohistochemistry

Paraffin embedded lung blocks from one donor whose medical record indicated a history and current medication usage of asthma whose cause of death was due to trauma underwent serial sectioning. Sections were stained with hematoxylin and eosin, anti-CD20 (1:500 dilution; Dako, Catalog Number M0755), anti-HLA-DQB1 (1:100 dilution; Atlas, Catalog Number HPA013667) or anti-HLA-DQB2 (1:500 dilution; Atlas, Catalog Number HPA073187).

## RESULTS

### Treatment-, timepoint- and cell-specific effects of innate and adaptive receptor stimulation on lung immune cells

We sought to determine the immune cell composition of human lungs and establish the gene expression responses following innate and adaptive immune cell activation at two different timepoints on gene expression. Immune cells from the UChicago Lung Biobank were used for these studies which includes leukocytes isolated from organ donors whose lungs were not used for transplantation(18). Cryopreserved cells in mixed leukocyte cell culture were separately treated with LPS, anti-CD3/CD28, anti-IgM/IgG + IL4 or media alone for 4- and 18-hours followed by scRNAseq.

Performing these experiments using the mixed leukocyte culture model enabled us to assess direct and indirect effects of treatments on cells (e.g. anti-CD3/CD28 on CD 4 T cells and anti-CD3/CD28 on B cells, respectively). As expected, most cells in our sample were macrophages and to enrich for lymphocytes, an additional step of cell sorting was performed on samples treated with anti-CD3/CD28 and anti-IgM/IgG prior to processing for scRNAseq. A total of 116,697 cells from six donors passed quality control and were used in our scRNAseq dataset. Cell annotation was able to classify macrophages into monocyte-derived and alveolar subsets and lymphocytes were broadly classified as B cells, CD4 T cells, CD8 T cells and NK/γδ cells (Figure 1A). We observed sub-clusters of cell populations that reflected treatment-by-timepoint effects on cell populations. For example, the majority of macrophages comprised a large cluster, but nearly all the macrophages treated with LPS for 4 hours could be separated into a distinct population that was also distinct from the macrophages treated with LPS for 18 hours (Figure 1B). Similarly, anti-IgM/IgG treated B cells could be identified as distinct from untreated B cells, and T cell subclusters were identified for distinct T cell subsets following anti-CD3/CD28 in the UMAP projection.

**Figure 1.**
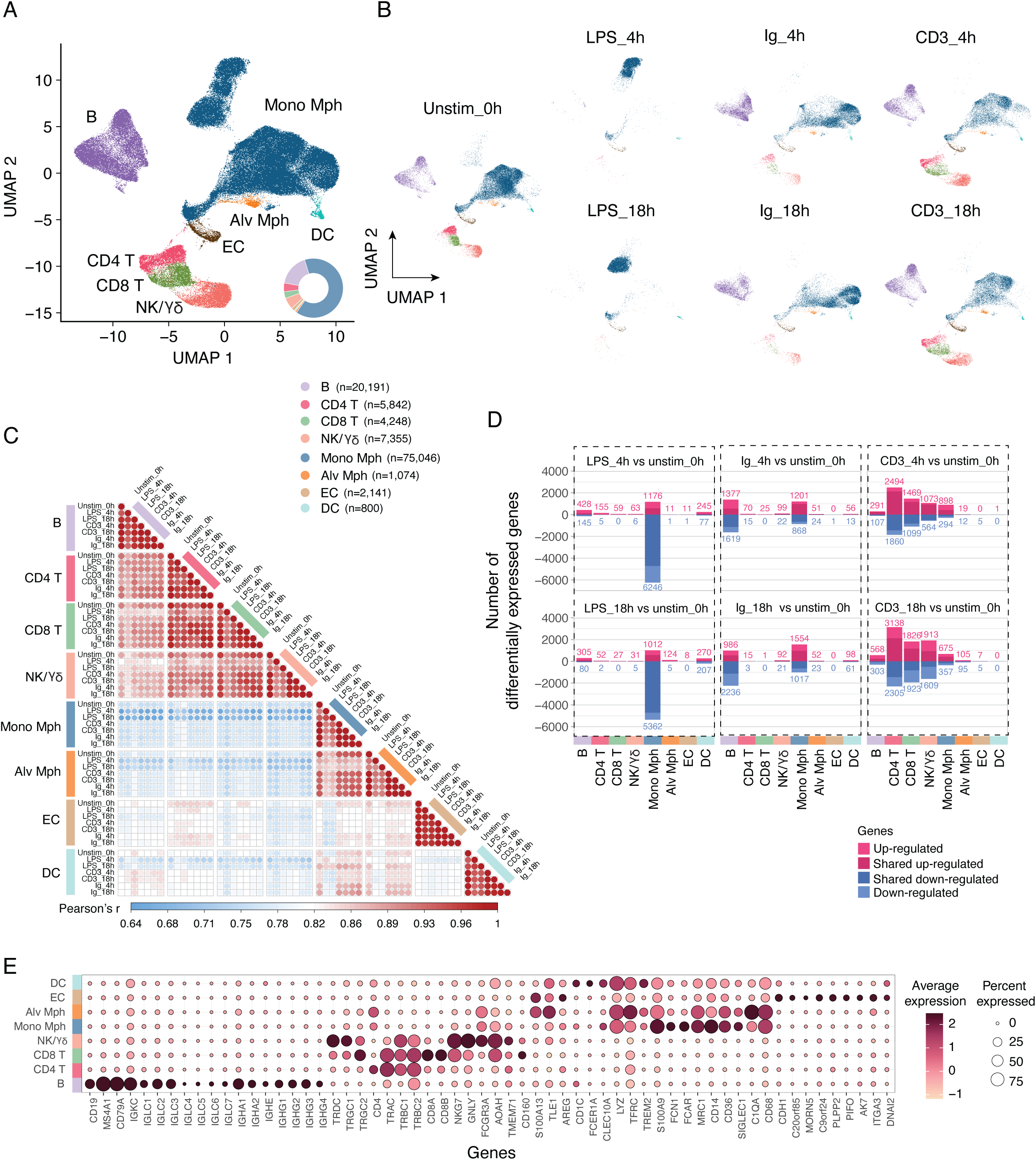
Cell clustering and annotations. A) UMAP visualization of annotated cell clusters colored by annotated cell type for all cells, treatments and timepoints. Inset indicates relative proportion of each cell type. Color key includes the number of cells annotated for each cell type. B) UMAP visualization of cells for each treatment and timepoint. C) Pearson’s correlation coefficient of gene expression between each cell type for each condition. D) Number of genes differentially expressed compared with the unstimulated control for each cell type plotted for each treatment and timepoint. Bars above and below 0 correspond to genes with higher expression and genes with lower expression, respectively. Area of bar plots with darker shading indicated genes differentially expressed for a cell type at both treatment timepoints. E) Average gene expression of selected cell marker genes. Alv Mph, alveolar macrophages; DC, dendritic cell; Mon Mph, monocyte-derived macrophage; NK/γδ, natural killer. CD3, anti-CD3/CD28 treatment; h, hours; Ig, anti-IgG/IgM treatment; LPS, lipopolysaccharide treatment; Unstim, unstimulated cells.

To assess the global gene expression patterns across cell types, we correlated gene expression for each cell type, treatment and timepoint. We observe relatively high correlation within a given cell type (Figure 1C). Correlation of gene expression was also higher within lymphocyte subsets (B cells, CD4 T cells, CD8 T cells and NK/γδ cells) for all treatments and timepoints than between lymphocyte subsets and the other cell types. Relatively high correlated gene expression was also observed between monocyte-derived macrophages and alveolar macrophages cells. Within both lung macrophage subsets, correlation of gene expression was reduced with LPS treatment, indicating the overall effects of TLR-4 stimulation on global gene expression. We then tested for gene expression differences between unstimulated and each treatment- and timepoint-condition and did this separately for each cell type, thus identifying the cell-specific effects of stimulation through different innate and adaptive immune receptors (Figure 1D). Treatment with LPS at both 4 hours and 18 hours induced DEGs across all cell types; the largest number of DEGs was in monocyte-derived macrophages with 7,422 genes differentially expressed at 4 hours and 6,374 genes differentially expressed at 18 hours and while the majority of these were differentially expressed at both timepoints. Timepoint-specific DEGs were also observed. Treatment with anti-IgM/IgG that targets the B cell receptor, resulted in 2,996 and 3,222 genes being differentially expressed in B cells at 4 and 18 hours, respectively. This treatment also induced robust DEGs in the monocyte-derived macrophage cells as well. With anti-CD3/CD28 treatment, CD4 T cells had the largest number of differentially expressed genes at both timepoints followed by CD8 T cells and NK/γδ cells. We also observed indirect effects on cell types with anti-CD3/CD28 treatment also resulting in differential expression of hundreds of genes in B cells at both timepoints.

To further evaluate gene expression of individual genes in each subset, we quantified the average expression and percent of each cell type that expressed a selected set of genes within a cell type for all treatment and timepoint conditions (Figure 1E). As expected, most B cells expressed *CD19*, *MS4A1*, and *CD79A*, and at high levels. Other genes expressed almost exclusively in B cells included *IGKC*, *IGLC* and *IGHC* genes that code for the B cell receptor proteins. As expected, T cell receptor genes *TRAC*, *TRBC1* and *TRBC2* were highly expressed in CD4 T cells, CD8 T cells and NK/γδ cells. Expression of the γδ chain genes, *TRGC1*, *TRGC2* and *TRDC* was present in the cells annotated as NK/γδ cells and some CD8 T cells.

### Lung immune cell responses are distinct from blood immune responses

We next sought to determine how similar lung immune cell responses were to blood immune cell responses. Our experiments were designed with similar treatments as in a prior study of blood immune cell populations(11). We therefore compared pseudo-bulk gene expression for the main cell types that responded to each treatment with the bulk RNA sequencing data from the Calderon et al study(11). Unstimulated lung B cell gene expression had the highest correlation with unstimulated blood naïve B cell gene expression (r = 0.54, = p-value 4.3×10^-308^) and had the lower correlation with gene expression in stimulated blood naïve, bulk and memory B cells (r = 0.38, p-value; r = 0.38, p-value = 7.5×10^-140^; and r = 6.5×10^-148^, respectively). For lung B cells at both anti-IgM/IgG treatment timepoints, correlation was also highest for unstimulated blood naïve B cells (r = 0.53, p-value = 4.9×10^-297^; and r = 0.54, p-value 5.4×10^-303^ at 4 hours and 18 hours, respectively), and correlation was also higher for unstimulated blood bulk and memory B cell populations than for any of the treated blood B cell populations (Figure 2).

**Figure 2.**
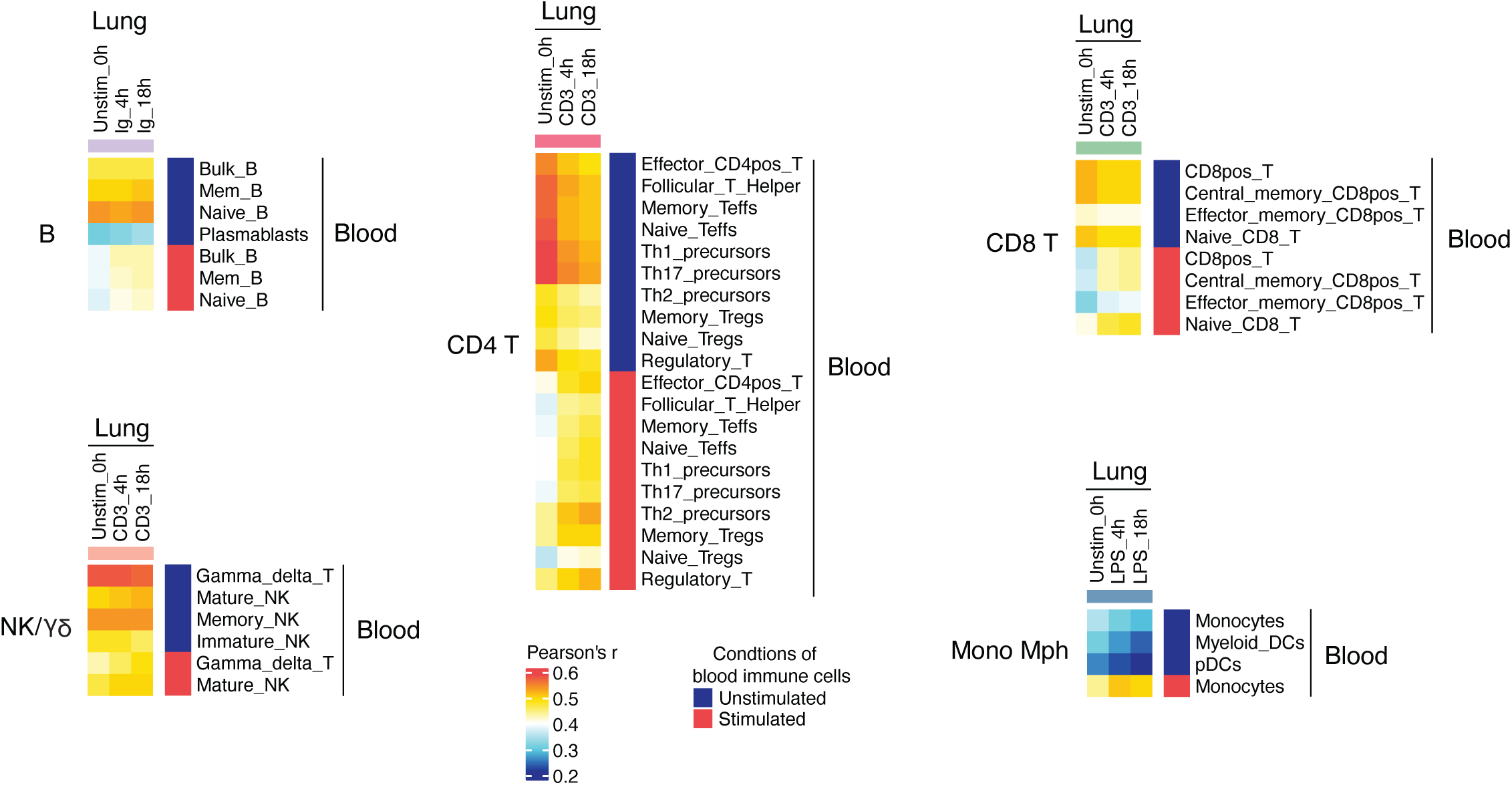
Gene expression correlations between lung and blood immune cell populations at resting and stimulated states. Gene lists of the union of differentially expressed genes at the 4 hour and 18 hour timepoints were generated for the main treatment targeting the annotated lung immune cell types (anti-IgM/IgG for B cells, anti-CD3/CD28 for NK/γδ and T cell subsets and LPS for macrophages) were generated. These gene lists were then used for the respective cell types to calculate correlation of expression between lung immune cells and expression in blood immune cells (Pearson’s correlation coefficient). Mon Mph, monocyte-derived macrophage.

**Figure 3.**
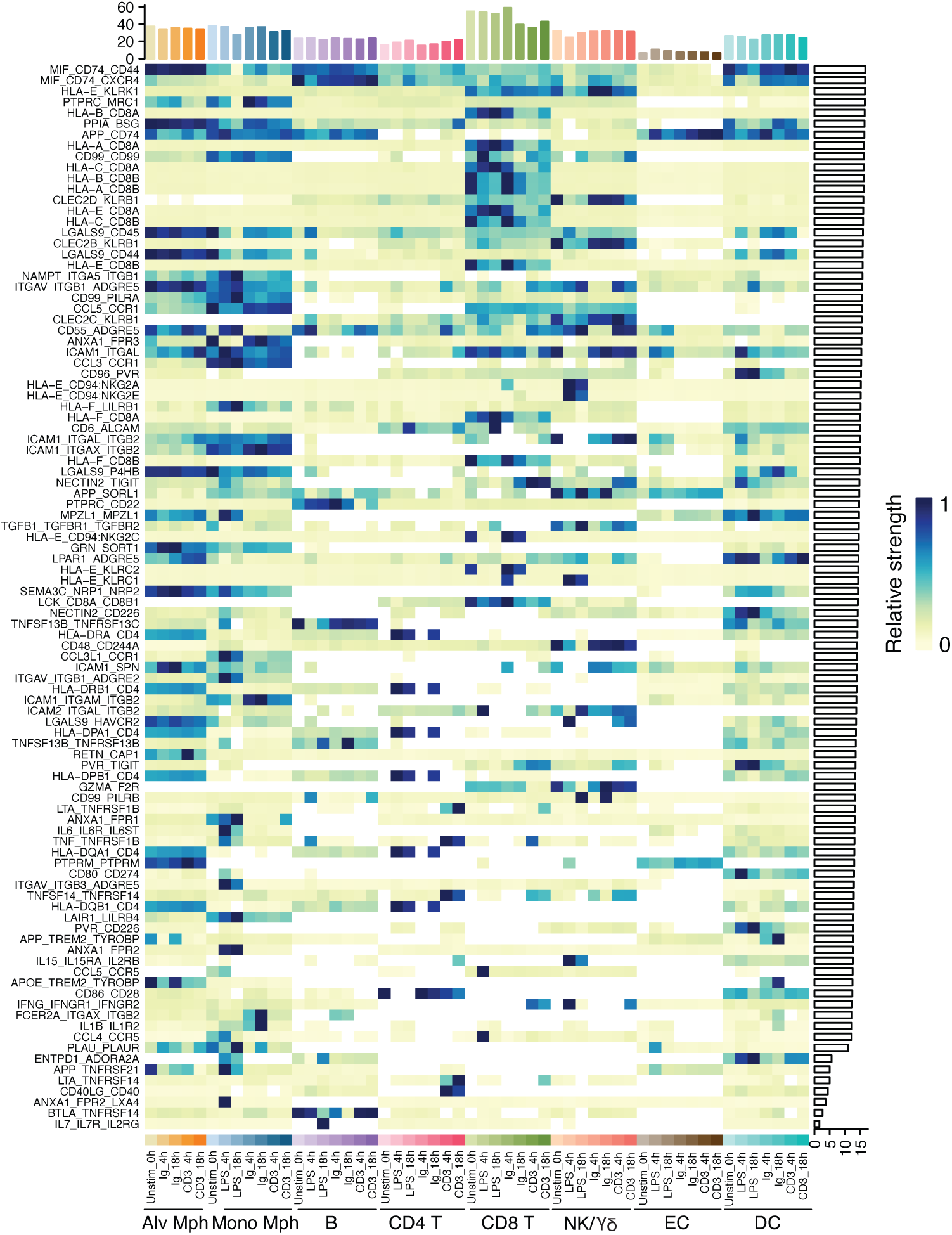
Inference of cell-to-cell contacts. Differentially expressed genes in any cell type following treatments were input to evaluate for potential receptor:ligand interactions. Analysis was conducted for each cell type stratified by treatment and timepoint and compared the relative strength of the probability of recptor:ligand interactions between a given cell type, condition and timepoint and all other cells in the dataset. All plotted receptor:ligand pairs are significant in at least one group of cells. Bars on top of figure indicate the relative strength of a cell group to overall receptor:ligand probabilities and horizontal bars on the right indicate that relative strength of a recptor:ligand pair across all cell groups.

For lung CD4 T cells, CD8 T cells and NK/γδ cells, we similarly observed the highest correlation for untreated and anti-CD3/CD28 treated cells at both timepoints was with cell subsets that had not been treated (Figure 2) and that overall, there was lower correlation of cell subsets after treatment. Macrophages were not included in the Calderon et al study, but blood monocytes, myeloid dendritic cells and plasmacytoid dendritic cells were included. Lung monocyte-derived macrophages had relatively low correlation with these three blood cells when unstimulated, but treatment of blood monocytes increased the correlation of gene expression for the lung macrophages (r = 0.44, p-value < 2.2×10^-308^; r = 0.51, p-value < 2.2×10^-308^; and r = 0.50, p-value < 2.2×10^-308^ for untreated, 4-hour LPS treated and 18-hour LPS treated, respectively). Collectively, these correlation results indicate that transcriptional responses for all studied lymphocyte populations are divergent between blood and lung cells.

### Lung immune cells dynamically alter receptors for cell-cell interactions

We evaluated the potential interactions between lung immune cells across all conditions by conducting receptor:ligand interaction analysis. This approach uses scRNAseq gene expression data to infer cellular interactions between a specified cell population and all other cells. To focus on receptor:ligand interactions that change with lung immune cell activation, we included any gene that was differentially expressed compared with the unstimulated state for all cell types. A total of 97 receptor:ligands interactions were identified as significant and 23 (23.7%) of these involved *HLA* genes (Figure 3A). Chemokine and cytokine signaling as well as costimulatory receptor:ligand interactions were also identified in this analysis. The macrophage migration inhibitory factor (MIF) pathway was involved in the top two receptor-ligand interactions (CD74:CD44 and CD74:CXCR4) and both of these differed in strength across cell types and treatments. CD8 T cells in all conditions had the strongest enrichment for receptor:ligand interactions, but many of these differed between treatments. For example, the HLA-B:CD8B was high in CD8 T cells that were not treated and at 4 hours after IgM/IgG treatment but was lower with LPS and anti-CDD3/CD28 treatments at both timepoints. This analysis demonstrates the plasticity of lung immune cells following stimulation of innate and adaptive immune receptors and effects broadly on interactions that mediate cell migration, cytokine and chemokine signaling, antigen presentation and costimulation.

### Lung immune cell programming across treatment conditions

To provide insights into cellular programs at different conditions, we separated the genes that were differentially expressed for each cell type’s main treatment response (anti-IgG/IgM for B cells, anti-CD3/CD28 for CD4 and CD8, and LPS for macrophages) based on whether they were increased or decreased and whether they were shared for both timepoints or specific to one timepoint and performed gene ontogeny enrichment on these lists of genes. We focused on the ‘Biological Processes’, ‘Molecular Functions’ and ‘Transcription Factors’ gene ontogeny sets. Overall, there were 7,862 gene ontogeny processes that were enriched in these four cell types. We observed enrichment in diverse cellular programs across the cell types. For B cells, genes with higher expression at both anti-IgM/IgG treatment timepoints were enriched in Biological Processes involved in lymphocyte activation; genes higher only at 4 hours of treatment included those involved in RNA processing while those higher only at 18 hours were enriched for cellular respiration (Figure 4A). Among the top Molecular Function enrichments for B cells, genes with higher expression at both timepoints included those involved in protein processing and antigen presentation; genes with higher expression only at 4 hours were enriched in RNA and protein binding while those with higher expression only at 18 hours were enriched for ATP-generating processes (Figure 4B).

**Figure 4.**
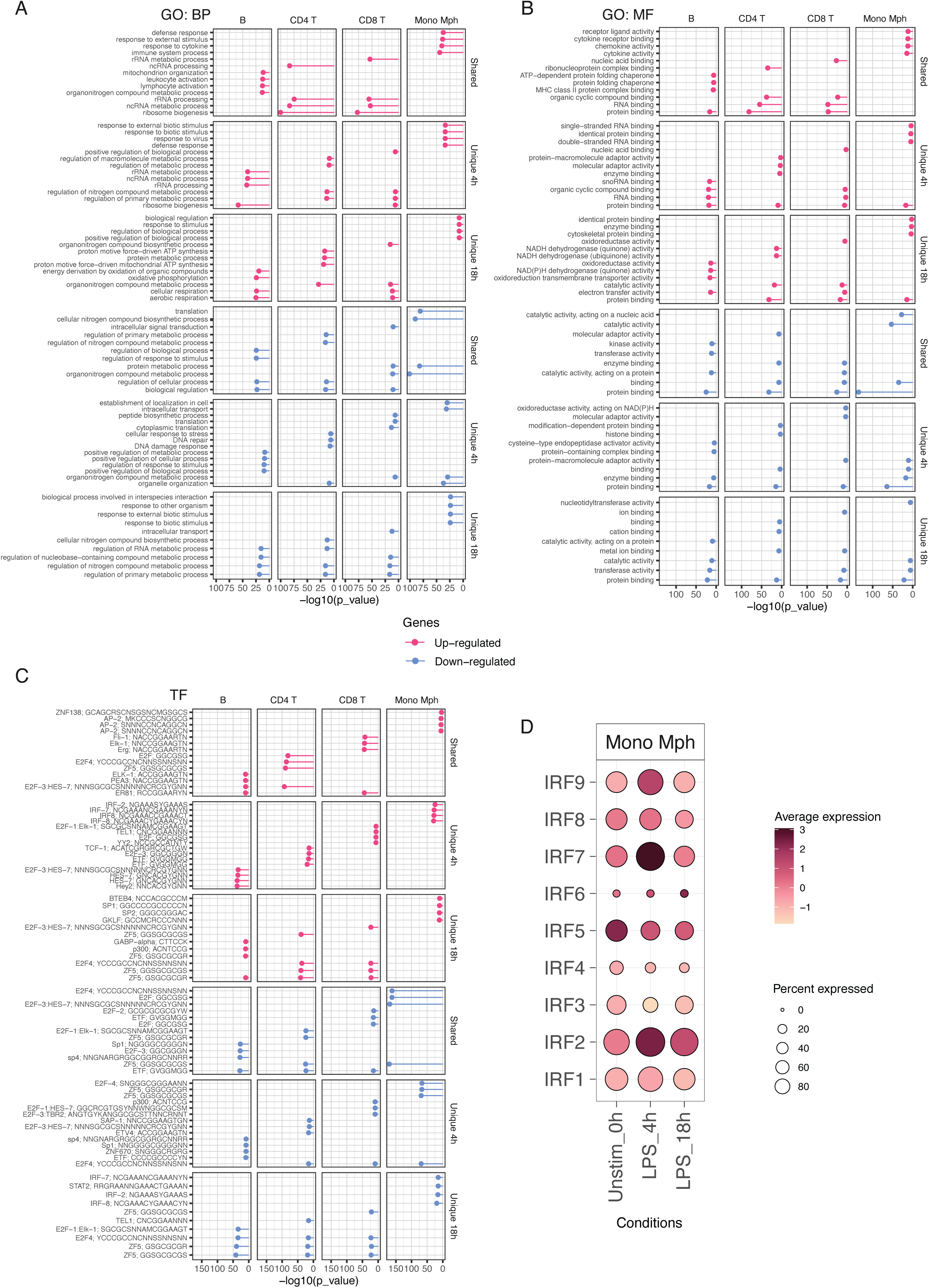
Top cellular programs identified through gene ontogeny analysis. Analyses were separately conducted for the main treatment targeting the annotated lung immune cell types (anti-IgM/IgG for B cells, anti-CD3/CD28 for T cell subsets and LPS for macrophages) using differentially expressed genes that were stratified by direction of effect and whether they were shared or unique to each timepoint. The top four gene ontogeny terms for each cell type are plotted for A) Biological Processes, B) Molecular Functions and C) Transcription Factors. D) Expression of *IRF* genes in monocyte-derived macrophages. BP, Biological Processes; MF, Molecular Functions; Mono Mph, monocyte-derived macrophage; TF, Transcription Factors.

The top gene ontogeny terms for genes that were more highly expressed at both timepoints for CD4 and CD8 T cells included genes involved in RNA processing (Biological Processing and Molecular Functions; Figures 4A-B). Interestingly, for CD4 and CD8 T cells, there was overlap of at least one top gene ontogeny term for 11/12 categories of gene sets in the Biological Processes and Molecular Functions, which indicates that despite these cell types have distinct responses following cellular activation the underlying cellular responses to anti-CD3/CD28 treatment are similar. To a lesser extent, there was also overlap terms between the T cells and the top gene ontogeny in B cells. The monocyte-derived macrophages had the least amount of sharing of gene ontogeny terms with other cell types.

We observed significant enrichment of genes with transcription factor motifs for all the gene lists (Figure 4C). In the monocyte-derived macrophages, genes that had increased expression at both timepoints following LPS treatment were enriched for genes with the transcription factor AP-2 motif binding sites. Interestingly, genes highly expressed only at 4 hours in monocyte-derived macrophages and genes that had lower expression at 18 hours were enriched for interferon response factor (IRF) transcription factors. These enrichments correlated with the expression of *IRF* genes that were differentially expressed in monocyte-derived macrophages: at 4 hours five *IRF* genes had increased expression (Log_2_ fold-change relative to unstimulated was 5.27 for *IRF1*, 8.07 for *IRF4*, 18.0 for *IRF7* and 3.18 for *IRF9*; Figure 4D) and at 18 hours four *IRF* genes had decreased expression (Log_2_ fold-change relative to unstimulated was −1.37 for *IRF1*, −1.33 for *IRF3*, −1.92 for *IRF5* and −1.74 for *IRF9*; Figure 4D). Our scRNAseq dataset reveals that there is dynamic gene regulation in monocyte-derived macrophages that is driven by early upregulation of *IRF* genes followed by reprogramming and downregulation of *IRF* genes and their targets following sustained LPS exposure.

### Differential expression of asthma-associated genes in activated lung immune cells

Previously, we performed a series of genome-wide association studies (GWASs) for childhood-onset asthma (COA) and adult-onset asthma (AOA) that identified shared and distinct genomic risk loci(14). Genetic variants and genes at these loci implicated immune cells as mediating a large portion of this genetic risk. We also observed multiple independent loci for COA and AOA at the HLA locus(16). We sought to further evaluate expression patterns of genes nominated as asthma-risk genes in our scRNAseq dataset to identify cell type- and treatment-related changes. We identified 225 genes at loci associated in either or both of our previous COA and AOA GWAS loci. For each cell type and condition, we tested for whether these genes were differentially expressed relative to the unstimulated state (Figure 5). Of these genes, 216 (96.0%) were differentially expressed in at least one cell type with treatment, and 197 (87.5%) of them were differentially expressed in more than one cell type or condition. We observed multiple distinct patterns of differential expression. For example, the shared COA and AOA gene *ERBB3* is differentially expressed in CD4 T cells at both timepoints following anti-CD3/CD28 treatment, but it is not differentially expressed in any other cell types or in CD4 T cells under any other conditions. Other genes, such as the COA gene *CCL20* is differentially expressed in many cell types (alveolar macrophages, B cells, CD4 T cells, endothelial cells, dendritic cells and monocyte-derived macrophages) but under different conditions. Our analysis of asthma-nominate genes in innate and adaptive immune cells in lungs shows that the overwhelming majority of these genes are differentially expressed in specific subsets following immune activation.

**Figure 5.**
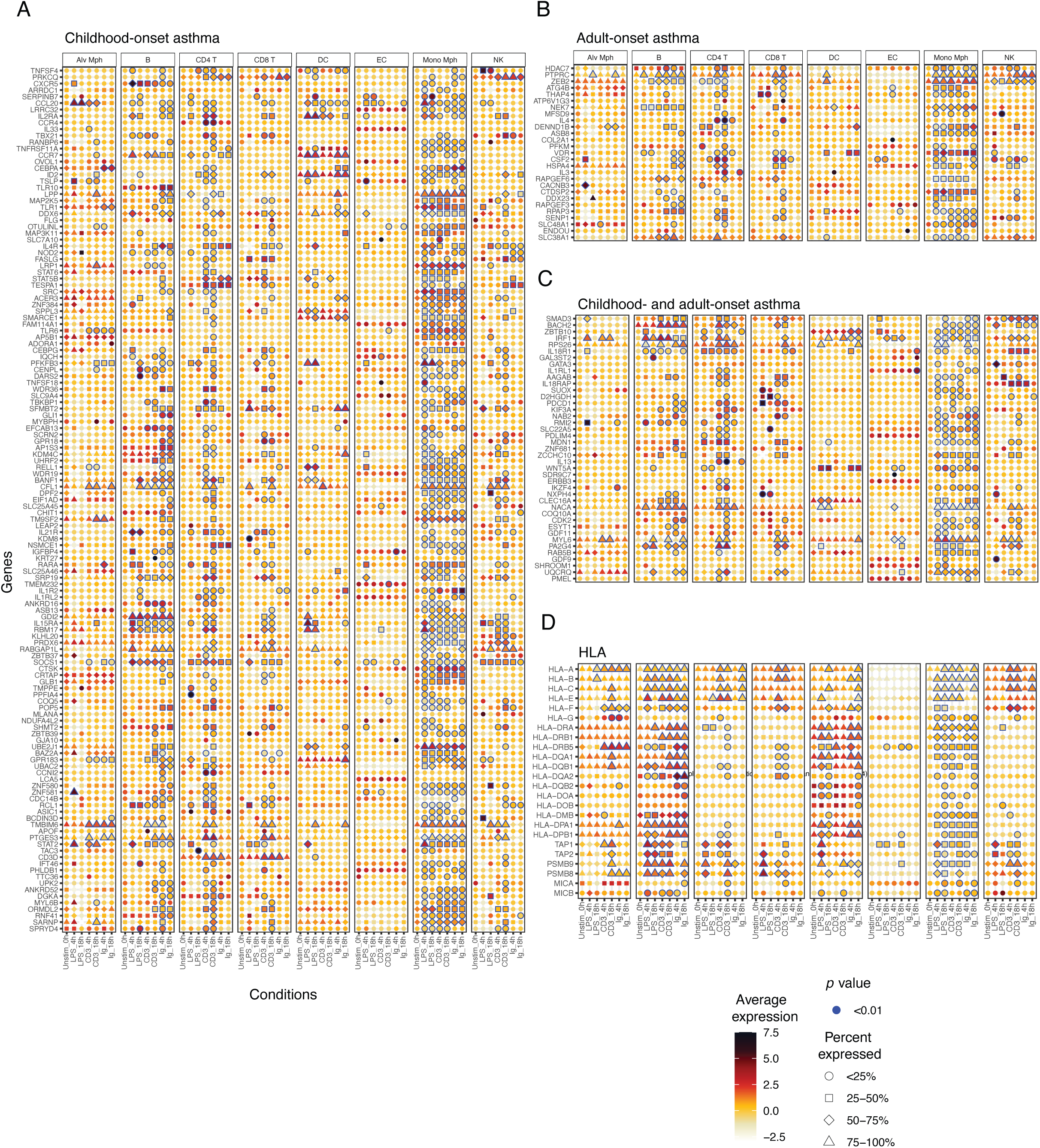
Expression patterns across lung immune cells of genes at loci associated with childhood-onset asthma and adult-onset asthma. A) Genes at COA loci. B) Genes at AOA loci. C) Genes at loci associated with COA and AOA. D) Genes at the HLA region. HLA, human leukocyte antigen.

**Figure 6.**
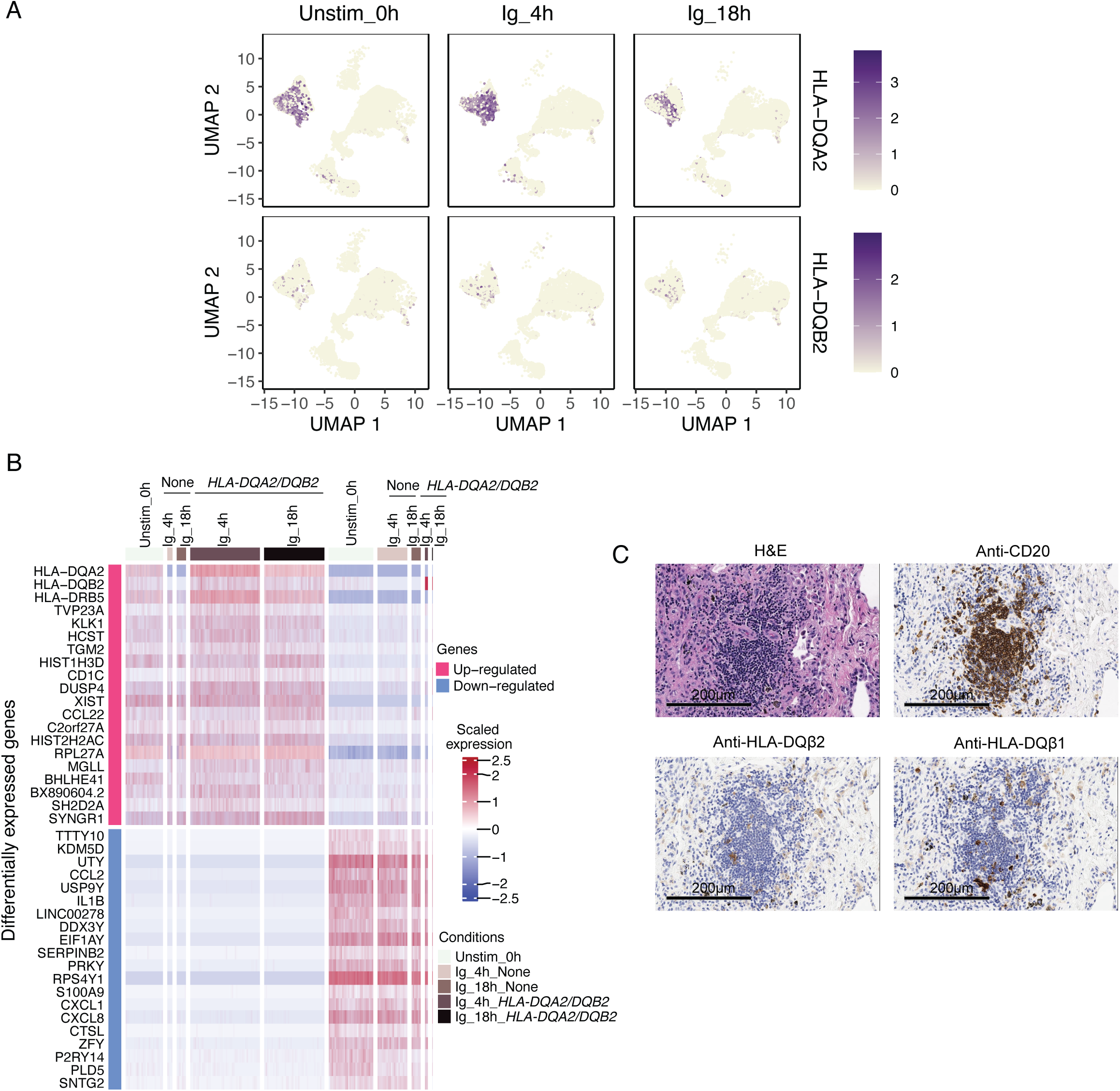
Asthma-associated and non-classical HLA class II genes *HLA-DQA2* and *HLA-DQB2* expression in lung B cells. A) UMAP plots in unstimulated and anti-IgM/IgG treatments colored for expression of *HLA-DQA2* and *HLA-DQB2*. B) Heatmap of the top 20 genes with increased and decreased expression in B cells that express one or both of these genes and B cells without detected expression of these genes. D) Hematoxylin and eosin staining and immunohistochemistry of serially sectioned slices of lung from organ donor with a history of asthma for CD20, HLA-DQβ1 and HLA-DQβ2. UMAP, uniform manifold approximation and projection.

### Expression of adult-onset asthma-associated non-classical HLA class II genes in lung B cells

In genetic and expression fine-mapping studies, we have previously nominated *HLA-DQA2* and *HLA-DQB2* as causal genes at the class II region for one of the AOA associations in this region(16). We therefore evaluated the expression of these genes in our scRNAseq dataset and identified B cells as the cell type with the highest percent and level of expression of these genes in unstimulated cells (Figure 5A). 4 hours after anti-IgM/IgG treatment, both *HLA-DQA2* and *HLA-DQB2* were more highly expressed in B cells (Log_2_-fold change 2.77 and 1.87; p-value_Adj_ 4.3×10^-205^ and 7.35×10^-14^, respectively) and expressed in a larger percentage of B cells (+27% and +6%, respectively). After 18 hours of treatment with anti-IgM/IgG, both *HLA-DQA2* and *HLA-DQB2* remained significantly increased compared with untreated B cells (Log_2_-fold change 2.03 and 1.28; p-value_Adj_ 6.18×10^-218^ and 5.58×10^-8^, respectively) and expressed in a larger percentage of B cells (+37% and +6%, respectively).

We then tested for gene expression differences between B cells that expressed *HLA-DQA2* and/or *HLA-DQB2* and those that do not, and 4,704 genes were differentially expressed. B cells that expressed these genes also had higher expression of *HLA-DRB5* as well as *CD1C* (Figure 5B). B cells without identified expression of either of these genes had higher expression of *IL1B*, *CCL2*, *CXCL1* and *CXCL8*. Finally, we sought to determine whether we could detect HLA-DQβ2, the protein product of *HLA-DQB2*) in human lungs. We performed immunohistochemistry on serially cut sections of lung from an organ donor whose clinical history included asthma and outpatient medications for treating asthma but whose cause of death was trauma. We identified bronchus-associated lymphoid tissue (BALT) within this donor that included a focus of CD20+ B cells (Figure 5C). A subset of cells in the BALT were positive for HLA-DQβ2.

## DISCUSSION

In this study we have leveraged the UChicago Lung Biobank resource to resolve human immune cell populations in the lung and interrogated dynamic responses of these populations following activation through TLR-4, B cell receptor and T cell receptor pathways. We characterized the main direct effects of LPS, anti-IgM/IgG and anti-CD3/CD28 on macrophages, B cells and T cells which are cell types with the highest expression of these respective receptor targets, and we identified indirect effects on lung immune cells that may be mediated by cytokine responses or cell-to-cell interactions. We demonstrate that correlation of gene expression between blood immune cells and lung immune cells is decreased in lymphocytes following treatments. We highlight changes with treatments in cell surface molecules that facilitate cell-to-cell interactions and the cellular processes for the main immune cell types. The expression of genes that have been nominated as asthma-relevant genes in genetic studies was characterized in lung immune cells and we observed large-scale changes in the expression of these genes with immune stimulation. Focusing on what we previously nominated as a the putatively causal genes through genetic fine mapping at the HLA class II region for AOA(16), we show that lung B cells are the immune cell type in the lung with the highest expression of *HLA-DQA2* and *HLA-DQB2* genes and that B cell receptor activation further induces upregulation of these genes.

A number of prior studies have characterized the immune cell populations present in human lungs and their gene expression profiles (2, 6, 7, 18), and some have focused on certain disease states such as asthma(1, 8), COPD(9) and viral infections(3, 5). One study included lung T cells and evaluated gene expression using a similar *in-vitro* stimulation with anti-CD3/CD28 and was able to catalog distinct T cell populations and states (21). Other studies have investigated the cellular responses following immune cell activation(11, 22–25). A common conclusion from these studies of immune cell responses is that there is dynamic regulation of gene expression that is context specific. Our study expands upon these in important ways. First, we investigated the effects of treatments that targeted innate and adaptive immune receptors. Second, we used a mixed leukocyte model with our stimulations which preserved the potential for cell-to-cell interactions which are crucial during *in vivo* immune responses. Third, we used cells isolated from the lungs, and our analyses raises concerns about the relevance of extrapolating observations made in blood immune cells to their counterparts in the lung. Finally, while we did not focus on contrasting the immune populations between donors with and without asthma, we were able to interrogate asthma-relevant genes identified in prior genetic studies.

We are interested in *HLA-DQA2* and *HLA-DQB2* because we have previously shown that AOA asthma risk genotype at the HLA class II locus is correlated with the expression of these genes in nasal epithelial cells, peripheral blood immune cells and lymphoblastoid cell lines (16). HLA class II genes encode proteins that combine as heterodimers to form the classical molecules HLA-DR (*HLA-DRA* and multiple HLA-DRB genes), HLA-DP (*HLA-DPA1* and *HLA-DPB1*) and HLA-DQ (*HLA-DQA1* and *HLA-DQB1*), and their expression is restricted to antigen presenting cells. These classical HLA class II molecules and genes are highly polymorphic and present peptides to CD4+ T cells, and most of the amino acid polymorphisms are in exon 2 at the peptide-binding region. Because these polymorphisms affect peptide binding, the overall effect is that they increase the diversity of ligands presented at the population level and therefore have a crucial role in the detection of pathogens and immunity. While the classical class II genes and their molecules have been extensively studied, surprising little is known about the AOA-associated *HLA-DQA2* and *HLA-DQB2* genes. *HLA-DQA2* and *HLA-DQB2* are paralogs of *HLA-DQA1* and *HLA-DQB1* but have a relative absence of polymorphisms(26), particularly in the peptide-binding domain. Interestingly, these genes are highly conserved in non-human primate species(27), but there is not a counterpart in mice. *HLA-DQA2* and *HLA-DQB2* were initially classified as pseudogenes, however, expression was ultimately identified in the skin and specifically in Langerhans cells(28), which are a specialized tissue antigen-presenting cell.

Lenormand et al. went on to detect the protein products of these genes, HLA-DQα2 and HLA-DQβ2, and showed they can form HLA-DQα2/HLA-DQβ2 heterodimers and traffic to the cell surface, as well as cross-heterodimers of HLA-DQα2 and HLA-DQβ1(28). In a study of the myeloproliferative disorder Langerhans cell histiocytosis, pathologic CD1a^+^CD207^+^ dendritic cells expressed *HLA-DQA2* and *HLA-DQB2* genes in bone marrow lesions(29). In a recent study, healthy adults were randomized to four weeks of treatment with inhaled corticosteroids (the most effective controller medication for asthma) or placebo and then underwent bronchoscopy with airway biopsy followed by bulk RNA sequencin(30). *HLA-DQA2* and *HLA-DQB2* were among the most downregulated genes by inhaled corticosteroids: expression was 1.36- and 1.51-fold lower in individuals who received treatment, respectively(30). Another study using a SARS-CoV-2 challenge of health adults showed that high expression of *HLA-DQA2* before inoculation was associated with preventing sustain infection(31).

We find that B cells expressing *HLA-DQA2* and/or *HLA-DQB2* also have increased expression of *CD1C* and *HLA-DRB5*. *CD1C* is related to major histocompatibility complex proteins and presents lipid and glycolipid self- and microbial-antigens to T cells and is expressed in dendritic cells, but has been reported to be expressed in B cells(32). Given the increased expression of these multiple classical and non-classical antigen presenting molecules, it is likely that B cells expressing *HLA-DQA2* or *HLA-DQB2* may be an inducible subset specialized for antigen presentation.

*CCL22* was also more highly expressed in these B cells and is also a potent chemoattractant for other immune cells. In contrast, B cells not expressing *HLA-DQA2* or *HLA-DQB2* expressed a different repertoire of chemoattractants, including *CXCL1* and *CXCL8*.

Our study has several limitations, which we acknowledge. While we flush blood vessels prior to isolation of leukocytes, it is possible that some of the lung immune cells included were transient populations. We included 4-hour and 18-hour timepoints in our experiments and it is likely that shorter or longer treatments would yield different results. While we used cells from 6 different donors, the number of cells that contributed to each cell type was not equal. We also included IL-4 with the anti-IgM/IgG treatment, so some of the gene expression changes could be related to this cytokine. We chose to broadly categorize cell types, and it is likely that sub-clustering further into additional cell types (e.g. naïve T cell, tissue resident memory T cells, effector memory, etc.) would generate different results. We did not have paired blood immune cells from donors and therefore relied on bulk RNA sequencing data generated in a previous study(11) for the comparison between lung and blood immune cell responses.

In conclusion, we observe cell type-, treatment- and timepoint-specific responses of lung innate and adaptive immune cells. Dynamic changes in gene expression were observed in all cell types, and the overwhelming majority of asthma-relevant genes exhibited differential expression in lung immune cells that was revealed with one or more types of stimulation. The non-classical class II genes *HLA-DQA2* and *HLA-DQB2* are expressed in lung antigen presenting cells and induced upon B cell activation the protein can be identified in lung BALT and may represent a novel antigen presentation pathway with relevance to asthma.

## References

1. Alladina J, Smith NP, Kooistra T, Slowikowski K, Kernin IJ, Deguine J, Keen HL, Manakongtreecheep K, Tantivit J, Rahimi RA, Sheng SL, Nguyen ND, Haring AM, Giacona FL, Hariri LP, Xavier RJ, Luster AD, Villani AC, Cho JL, Medoff BD. A human model of asthma exacerbation reveals transcriptional programs and cell circuits specific to allergic asthma. Sci Immunol 2023; 8: eabq6352.

2. Dominguez Conde C, Xu C, Jarvis LB, Rainbow DB, Wells SB, Gomes T, Howlett SK, Suchanek O, Polanski K, King HW, Mamanova L, Huang N, Szabo PA, Richardson L, Bolt L, Fasouli ES, Mahbubani KT, Prete M, Tuck L, Richoz N, Tuong ZK, Campos L, Mousa HS, Needham EJ, Pritchard S, Li T, Elmentaite R, Park J, Rahmani E, Chen D, Menon DK, Bayraktar OA, James LK, Meyer KB, Yosef N, Clatworthy MR, Sims PA, Farber DL, Saeb-Parsy K, Jones JL, Teichmann SA. Cross-tissue immune cell analysis reveals tissue-specific features in humans. Science 2022; 376: eabl5197.

3. Grant RA, Morales-Nebreda L, Markov NS, Swaminathan S, Querrey M, Guzman ER, Abbott DA, Donnelly HK, Donayre A, Goldberg IA, Klug ZM, Borkowski N, Lu Z, Kihshen H, Politanska Y, Sichizya L, Kang M, Shilatifard A, Qi C, Lomasney JW, Argento AC, Kruser JM, Malsin ES, Pickens CO, Smith SB, Walter JM, Pawlowski AE, Schneider D, Nannapaneni P, Abdala-Valencia H, Bharat A, Gottardi CJ, Budinger GRS, Misharin AV, Singer BD, Wunderink RG, Investigators NSS. Circuits between infected macrophages and T cells in SARS-CoV-2 pneumonia. Nature 2021; 590: 635–641.

4. Raudvere U, Kolberg L, Kuzmin I, Arak T, Adler P, Peterson H, Vilo J. g:Profiler: a web server for functional enrichment analysis and conversions of gene lists (2019 update). Nucleic Acids Res 2019; 47: W191–W198.

5. Ren X, Wen W, Fan X, Hou W, Su B, Cai P, Li J, Liu Y, Tang F, Zhang F, Yang Y, He J, Ma W, He J, Wang P, Cao Q, Chen F, Chen Y, Cheng X, Deng G, Deng X, Ding W, Feng Y, Gan R, Guo C, Guo W, He S, Jiang C, Liang J, Li YM, Lin J, Ling Y, Liu H, Liu J, Liu N, Liu SQ, Luo M, Ma Q, Song Q, Sun W, Wang G, Wang F, Wang Y, Wen X, Wu Q, Xu G, Xie X, Xiong X, Xing X, Xu H, Yin C, Yu D, Yu K, Yuan J, Zhang B, Zhang P, Zhang T, Zhao J, Zhao P, Zhou J, Zhou W, Zhong S, Zhong X, Zhang S, Zhu L, Zhu P, Zou B, Zou J, Zuo Z, Bai F, Huang X, Zhou P, Jiang Q, Huang Z, Bei JX, Wei L, Bian XW, Liu X, Cheng T, Li X, Zhao P, Wang FS, Wang H, Su B, Zhang Z, Qu K, Wang X, Chen J, Jin R, Zhang Z. COVID-19 immune features revealed by a large-scale single-cell transcriptome atlas. Cell 2021; 184: 5838.

6. Shilts J, Severin Y, Galaway F, Muller-Sienerth N, Chong ZS, Pritchard S, Teichmann S, Vento-Tormo R, Snijder B, Wright GJ. A physical wiring diagram for the human immune system. Nature 2022; 608: 397–404.

7. Travaglini KJ, Nabhan AN, Penland L, Sinha R, Gillich A, Sit RV, Chang S, Conley SD, Mori Y, Seita J, Berry GJ, Shrager JB, Metzger RJ, Kuo CS, Neff N, Weissman IL, Quake SR, Krasnow MA. A molecular cell atlas of the human lung from single-cell RNA sequencing. Nature 2020; 587: 619–625.

8. Vieira Braga FA, Kar G, Berg M, Carpaij OA, Polanski K, Simon LM, Brouwer S, Gomes T, Hesse L, Jiang J, Fasouli ES, Efremova M, Vento-Tormo R, Talavera-Lopez C, Jonker MR, Affleck K, Palit S, Strzelecka PM, Firth HV, Mahbubani KT, Cvejic A, Meyer KB, Saeb-Parsy K, Luinge M, Brandsma CA, Timens W, Angelidis I, Strunz M, Koppelman GH, van Oosterhout AJ, Schiller HB, Theis FJ, van den Berge M, Nawijn MC, Teichmann SA. A cellular census of human lungs identifies novel cell states in health and in asthma. Nat Med 2019; 25: 1153–1163.

9. Villasenor-Altamirano AB, Jain D, Jeong Y, Menon JA, Kamiya M, Haider H, Manandhar R, Sheikh MDA, Athar H, Merriam LT, Ryu MH, Sasaki T, Castaldi PJ, Rao DA, Sholl LM, Vivero M, Hersh CP, Zhou X, Veerkamp J, Yun JH, Kim EY, MassGeneralBrigham - Bayer Pulmonary Drug Discovery L. Activation of CD8(+) T Cells in Chronic Obstructive Pulmonary Disease Lung. Am J Respir Crit Care Med 2023; 208: 1177–1195.

10. Schmiedel BJ, Gonzalez-Colin C, Fajardo V, Rocha J, Madrigal A, Ramirez-Suastegui C, Bhattacharyya S, Simon H, Greenbaum JA, Peters B, Seumois G, Ay F, Chandra V, Vijayanand P. Single-cell eQTL analysis of activated T cell subsets reveals activation and cell type-dependent effects of disease-risk variants. Sci Immunol 2022; 7: eabm2508.

11. Calderon D, Nguyen MLT, Mezger A, Kathiria A, Muller F, Nguyen V, Lescano N, Wu B, Trombetta J, Ribado JV, Knowles DA, Gao Z, Blaeschke F, Parent AV, Burt TD, Anderson MS, Criswell LA, Greenleaf WJ, Marson A, Pritchard JK. Landscape of stimulation-responsive chromatin across diverse human immune cells. Nat Genet 2019; 51: 1494–1505.

12. Demenais F, Margaritte-Jeannin P, Barnes KC, Cookson WOC, Altmuller J, Ang W, Barr RG, Beaty TH, Becker AB, Beilby J, Bisgaard H, Bjornsdottir US, Bleecker E, Bonnelykke K, Boomsma DI, Bouzigon E, Brightling CE, Brossard M, Brusselle GG, Burchard E, Burkart KM, Bush A, Chan-Yeung M, Chung KF, Couto Alves A, Curtin JA, Custovic A, Daley D, de Jongste JC, Del-Rio-Navarro BE, Donohue KM, Duijts L, Eng C, Eriksson JG, Farrall M, Fedorova Y, Feenstra B, Ferreira MA, Australian Asthma Genetics Consortium c, Freidin MB, Gajdos Z, Gauderman J, Gehring U, Geller F, Genuneit J, Gharib SA, Gilliland F, Granell R, Graves PE, Gudbjartsson DF, Haahtela T, Heckbert SR, Heederik D, Heinrich J, Heliovaara M, Henderson J, Himes BE, Hirose H, Hirschhorn JN, Hofman A, Holt P, Hottenga J, Hudson TJ, Hui J, Imboden M, Ivanov V, Jaddoe VWV, James A, Janson C, Jarvelin MR, Jarvis D, Jones G, Jonsdottir I, Jousilahti P, Kabesch M, Kahonen M, Kantor DB, Karunas AS, Khusnutdinova E, Koppelman GH, Kozyrskyj AL, Kreiner E, Kubo M, Kumar R, Kumar A, Kuokkanen M, Lahousse L, Laitinen T, Laprise C, Lathrop M, Lau S, Lee YA, Lehtimaki T, Letort S, Levin AM, Li G, Liang L, Loehr LR, London SJ, Loth DW, Manichaikul A, Marenholz I, Martinez FJ, Matheson MC, Mathias RA, Matsumoto K, Mbarek H, McArdle WL, Melbye M, Melen E, Meyers D, Michel S, Mohamdi H, Musk AW, Myers RA, Nieuwenhuis MAE, Noguchi E, O’Connor GT, Ogorodova LM, Palmer CD, Palotie A, Park JE, Pennell CE, Pershagen G, Polonikov A, Postma DS, Probst-Hensch N, Puzyrev VP, Raby BA, Raitakari OT, Ramasamy A, Rich SS, Robertson CF, Romieu I, Salam MT, Salomaa V, Schlunssen V, Scott R, Selivanova PA, Sigsgaard T, Simpson A, Siroux V, Smith LJ, Solodilova M, Standl M, Stefansson K, Strachan DP, Stricker BH, Takahashi A, Thompson PJ, Thorleifsson G, Thorsteinsdottir U, Tiesler CMT, Torgerson DG, Tsunoda T, Uitterlinden AG, van der Valk RJP, Vaysse A, Vedantam S, von Berg A, von Mutius E, Vonk JM, Waage J, Wareham NJ, Weiss ST, White WB, Wickman M, Widen E, Willemsen G, Williams LK, Wouters IM, Yang JJ, Zhao JH, Moffatt MF, Ober C, Nicolae DL. Multiancestry association study identifies new asthma risk loci that colocalize with immune-cell enhancer marks. Nat Genet 2018; 50: 42–53.

13. Ferreira MAR, Mathur R, Vonk JM, Szwajda A, Brumpton B, Granell R, Brew BK, Ullemar V, Lu Y, Jiang Y, and Me Research T, e QC, Consortium B, Magnusson PKE, Karlsson R, Hinds DA, Paternoster L, Koppelman GH, Almqvist C. Genetic Architectures of Childhood- and Adult-Onset Asthma Are Partly Distinct. Am J Hum Genet 2019; 104: 665–684.

14. Pividori M, Schoettler N, Nicolae DL, Ober C, Im HK. Shared and distinct genetic risk factors for childhood-onset and adult-onset asthma: genome-wide and transcriptome-wide studies. Lancet Respir Med 2019; 7: 509–522.

15. Tsuo K, Zhou W, Wang Y, Kanai M, Namba S, Gupta R, Majara L, Nkambule LL, Morisaki T, Okada Y, Neale BM, Global Biobank Meta-analysis I, Daly MJ, Martin AR. Multi-ancestry meta-analysis of asthma identifies novel associations and highlights the value of increased power and diversity. Cell Genom 2022; 2: 100212.

16. Clay SM, Schoettler N, Goldstein AM, Carbonetto P, Dapas M, Altman MC, Rosasco MG, Gern JE, Jackson DJ, Im HK, Stephens M, Nicolae DL, Ober C. Fine-mapping studies distinguish genetic risks for childhood- and adult-onset asthma in the HLA region. Genome Med 2022; 14: 55.

17. Hrusch CL, Manns ST, Bryazka D, Casaos J, Bonham CA, Jaffery MR, Blaine KM, Mills KAM, Verhoef PA, Adegunsoye AO, Williams JW, Tjota MY, Moore TV, Strek ME, Noth I, Sperling AI. ICOS protects against mortality from acute lung injury through activation of IL-5(+) ILC2s. Mucosal Immunol 2018; 11: 61–70.

18. Schoettler N, Hrusch CL, Blaine KM, Sperling AI, Ober C. Transcriptional programming and T cell receptor repertoires distinguish human lung and lymph node memory T cells. Commun Biol 2019; 2: 411.

19. Stuart T, Butler A, Hoffman P, Hafemeister C, Papalexi E, Mauck WM3rd, Hao Y, Stoeckius M, Smibert P, Satija R. Comprehensive Integration of Single-Cell Data. Cell 2019; 177: 1888–1902 e1821.

20. DePasquale EAK, Schnell DJ, Van Camp PJ, Valiente-Alandi I, Blaxall BC, Grimes HL, Singh H, Salomonis N. DoubletDecon: Deconvoluting Doublets from Single-Cell RNA-Sequencing Data. Cell Rep 2019; 29: 1718–1727 e1718.

21. Szabo PA, Levitin HM, Miron M, Snyder ME, Senda T, Yuan J, Cheng YL, Bush EC, Dogra P, Thapa P, Farber DL, Sims PA. Single-cell transcriptomics of human T cells reveals tissue and activation signatures in health and disease. Nat Commun 2019; 10: 4706.

22. Amarasinghe HE, Zhang P, Whalley JP, Allcock A, Migliorini G, Brown AC, Scozzafava G, Knight JC. Mapping the epigenomic landscape of human monocytes following innate immune activation reveals context-specific mechanisms driving endotoxin tolerance. BMC Genomics 2023; 24: 595.

23. Zhang P, Amarasinghe HE, Whalley JP, Tay C, Fang H, Migliorini G, Brown AC, Allcock A, Scozzafava G, Rath P, Davies B, Knight JC. Epigenomic analysis reveals a dynamic and context-specific macrophage enhancer landscape associated with innate immune activation and tolerance. Genome Biol 2022; 23: 136.

24. Soskic B, Cano-Gamez E, Smyth DJ, Ambridge K, Ke Z, Matte JC, Bossini-Castillo L, Kaplanis J, Ramirez-Navarro L, Lorenc A, Nakic N, Esparza-Gordillo J, Rowan W, Wille D, Tough DF, Bronson PG, Trynka G. Immune disease risk variants regulate gene expression dynamics during CD4(+) T cell activation. Nat Genet 2022; 54: 817–826.

25. Bediaga NG, Coughlan HD, Johanson TM, Garnham AL, Naselli G, Schroder J, Fearnley LG, Bandala-Sanchez E, Allan RS, Smyth GK, Harrison LC. Multi-level remodelling of chromatin underlying activation of human T cells. Sci Rep 2021; 11: 528.

26. Berdoz J, Tiercy JM, Rollini P, Mach B, Gorski J. Remarkable sequence conservation of the HLA-DQB2 locus (DX beta) within the highly polymorphic DQ subregion of the human MHC. Immunogenetics 1989; 29: 241–248.

27. Gaur LK, Heise ER, Thurtle PS, Nepom GT. Conservation of the HLA-DQB2 locus in nonhuman primates. J Immunol 1992; 148: 943–948.

28. Lenormand C, Bausinger H, Gross F, Signorino-Gelo F, Koch S, Peressin M, Fricker D, Cazenave JP, Bieber T, Hanau D, de la Salle H, Tourne S. HLA-DQA2 and HLA-DQB2 genes are specifically expressed in human Langerhans cells and encode a new HLA class II molecule. J Immunol 2012; 188: 3903–3911.

29. Lim KPH, Milne P, Poidinger M, Duan K, Lin H, McGovern N, Abhyankar H, Zinn D, Burke TM, Eckstein OS, Chakraborty R, Sengal A, Scull B, Newell E, Merad M, McClain KL, Man TK, Ginhoux F, Collin M, Allen CE. Circulating CD1c+ myeloid dendritic cells are potential precursors to LCH lesion CD1a+CD207+ cells. Blood Adv 2020; 4: 87–99.

30. Marchi E, Hinks TSC, Richardson M, Khalfaoui L, Symon FA, Rajasekar P, Clifford R, Hargadon B, Austin CD, MacIsaac JL, Kobor MS, Siddiqui S, Mar JS, Arron JR, Choy DF, Bradding P. The effects of inhaled corticosteroids on healthy airways. Allergy 2024; 79: 1831–1843.

31. Lindeboom RGH, Worlock KB, Dratva LM, Yoshida M, Scobie D, Wagstaffe HR, Richardson L, Wilbrey-Clark A, Barnes JL, Kretschmer L, Polanski K, Allen-Hyttinen J, Mehta P, Sumanaweera D, Boccacino JM, Sungnak W, Elmentaite R, Huang N, Mamanova L, Kapuge R, Bolt L, Prigmore E, Killingley B, Kalinova M, Mayer M, Boyers A, Mann A, Swadling L, Woodall MNJ, Ellis S, Smith CM, Teixeira VH, Janes SM, Chambers RC, Haniffa M, Catchpole A, Heyderman R, Noursadeghi M, Chain B, Mayer A, Meyer KB, Chiu C, Nikolic MZ, Teichmann SA. Human SARS-CoV-2 challenge uncovers local and systemic response dynamics. Nature 2024; 631: 189–198.

32. Allan LL, Stax AM, Zheng DJ, Chung BK, Kozak FK, Tan R, van den Elzen P. CD1d and CD1c expression in human B cells is regulated by activation and retinoic acid receptor signaling. J Immunol 2011; 186: 5261–5272.

